# In silico investigation of biomechanical response of a human subjected to primary blast

**DOI:** 10.1101/2021.09.16.460591

**Authors:** Sunil Sutar, S. G. Ganpule

## Abstract

The response of the brain to the explosion induced primary blast waves is actively sought. Over the past decade, reasonable progress has been made in the fundamental understanding of bTBI using head surrogates and animal models. Yet, the current understanding of how blast waves interact with the human is in nascent stages, primarily due to lack of data in humans. The biomechanical response in human is critically required so that connection to the aforementioned bTBI models can be faithfully established. Here, using a detailed, full-body human model, we elucidate the biomechanical cascade of the brain under a primary blast. The input to the model is incident overpressure as achieved by specifying charge mass and standoff distance through ConWep. The full-body model allows to holistically probe short- (<5 ms) and long-term (200 ms) brain biomechanical responses. The full-body model has been extensively validated against impact loading in the past. In this work, we validate the head model against blast loading. We also incorporate structural anisotropy of the brain white matter. Blast wave human interaction is modeled using a conventional weapon modeling approach. We demonstrate that the blast wave transmission, linear and rotational motion of the head are dominant pathways for the biomechanical loading of the brain, and these loading paradigms generate distinct biomechanical fields within the brain. Blast transmission and linear motion of the head govern the volumetric response, whereas the rotational motion of the head governs the deviatoric response. We also observe that blast induced head rotation alone produces a diffuse injury pattern in white matter fiber tracts. Lastly, we find that the biomechanical response under blast is comparable to the impact event. These insights will augment laboratory and clinical investigations of bTBI and help devise better blast mitigation strategies.

## 1. Introduction

Blast induced traumatic brain injury (bTBI) is a substantial health concern in the era of asymmetric warfare.(Tanielian, 2008; Warden, 2006) Understanding how the blast wave interacts with the head-brain parenchyma is crucial for devising blast mitigation strategies, clinical interventions, therapeutics, and recreating injury in a laboratory. Several mechanisms of blast induced traumatic brain injury have been proposed, including direct intracranial transmission,(Nyein et al., 2010; Sundaramurthy et al., 2012; Taylor and Ford, 2009) skull flexure,(Bolander et al., 2011; Garimella et al., 2018; Moss et al., 2009) a thoracic mechanism,(Cernak et al., 2001; Simard et al., 2014) cavitation,(Panzer et al., 2012; Salzar et al., 2017) and head acceleration.(Goldstein et al., 2012; Gullotti et al., 2014) Yet, the current understanding of the contribution of each mechanism in generating mechanical fields within the brain is limited (Fievisohn et al., 2018; Meaney et al., 2014) and findings in the literature are contradictory.(Fievisohn et al., 2018) The confounding findings in the literature are mainly attributed to various factors, including differences in experimental head models, blast wave generator, resulting blast characteristic, and lack of standardization in biomechanical measurements. Experiments on living humans and primates are unfeasible due to ethical reasons. On the other hand, clinical investigations of the effects of the pure primary blast, with the absence of secondary and tertiary effects,(Ling et al., 2009) are limited. In silico investigations using computational models offer an attractive alternative to study the effects of blast, provided computational models are robust enough to capture the physics.

Blast injury is classified as primary, secondary, tertiary, and quaternary injury depending on the external mechanical insult. Primary blast injury results from the impact of the explosion induced blast wave with body surfaces. The secondary injury occurs from matter thrown by the explosion (e.g., fragments, flying debris). Tertiary injury results when the body or part of body is thrown by the blast wind and strikes intervening objects such as a wall or the ground. Quaternary injury includes all other explosion related injuries (e.g., injury due to the burns).

In this work, using a full-body human model (Fig. 1), we examine the spatiotemporal evolution of the mechanical fields within the brain under the primary blast as the blast interacts with the body. We have only simulated the effects of explosion induced blast wave and hence investigated a primary blast. The input to the model is incident (static) overpressure as achieved by specifying charge mass and standoff distance through ConWep (see Supplementary Fig. S1 and Methods section for details). Compared to earlier investigations, we specifically investigate the response of full-body human (as opposed to only head) for longer durations (200 ms). The simulated length- and time-scales allow to probe the holistic response of human against the primary blast. Using carefully constructed strategies, we also delineate the effects of wave transmission and global head motion on mechanical fields within the human brain.

**Figure 1:**
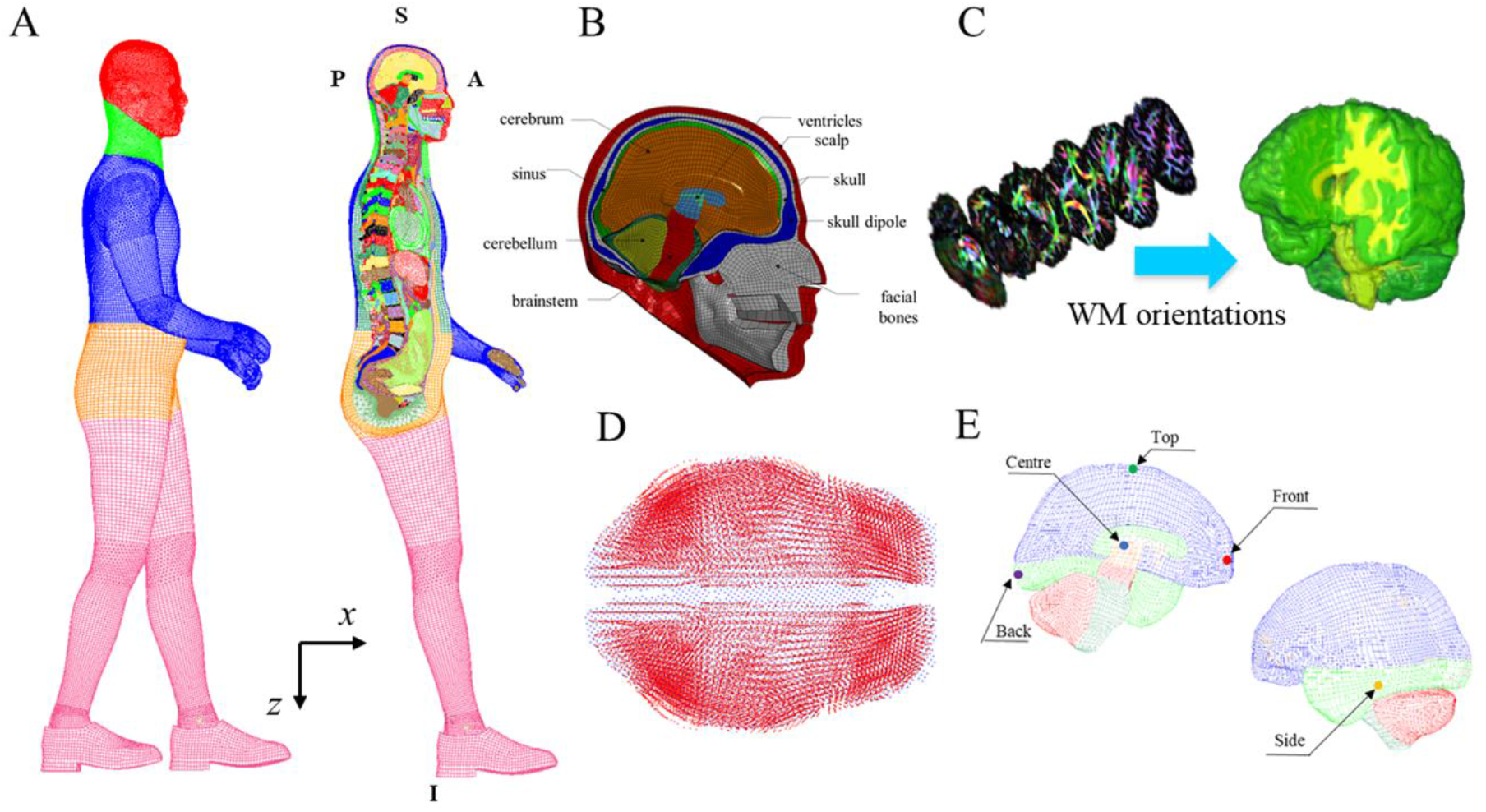
Full-body, computational human model in standing position. (A) The model is broadly divided into five body regions: head (red), neck (green), thorax (blue), abdomen (orange), and pelvic-lower extremity (magenta). (B) Head Model consists of the scalp, skull, facial bones, sinuses, dura, pia, arachnoid, falx, tentorium, bridging veins, and brain. (C) The white matter fiber orientations are incorporated into the existing head model using diffusion tensor imaging. (D) DTI data (blue) was aligned with the finite element model (red) using rigid body translation and rotation. (E) Representative locations in the brain those are used to probe time history plots.

## 2. Methods

### 2.1 Full-body Human Model

The full-body, computational human model in standing position used in this work (Fig. 1A) is based on Global Human Body Models Consortium, GHBMC.(Gayzik et al., 2011a; Gayzik et al., 2009; Gayzik et al., 2011b) The full-body human model, including the head model (Fig. 1B),(Mao et al., 2013) is extensively validated (Supplementary Table S1) under blunt impact loading through multi-institutional effort. In this work, we make the following contributions to the head model: (a) We extend the applicability of the head model by validating it against a range of blast scenarios. We also compare model response against experimentally measured *in vivo* strains in human volunteers. (b) We incorporate fiber orientations of the white matter and compute strain along the fiber direction. (c) In addition to the baseline constitutive model, we incorporate the material model and associated high strain rate properties for the brain tissue that are relevant for the blast loading.

We purposefully chose full-body model for a few particular reasons. First, we wanted to simulate the response of the whole human as opposed to the only head. Second, we wanted to use biofidelic neck (as implemented in GHBMC, details in Supplementary Information) that plays a crucial role in faithfully capturing head kinematics in response to the blast, especially at longer times. The important details of the full-body model (especially, head and neck model) are provided in Supplementary Information.

### 2.2 Loading and Boundary Conditions

The interaction of blast wave with the full-body human model was simulated using the conventional weapon (ConWep) modeling approach.(Hyde, 2004; LSTC, 2017) ConWep simulates the propagation of free-field blast waves in an infinite domain. The incident and reflected pressures at various locations of the target are computed based on explosive mass and stand-off distance using empirical relations.(Kingery and Bulmash, 1984)

We have considered three blast intensities with peak incident overpressures of ∼70, ∼140, ∼210 kPa, and a positive phase duration of ∼6 ms (Supplementary Fig. S1); which are relevant to bTBI injury.(Rafaels et al., 2012) The results of 210 kPa blast intensity are presented, unless stated otherwise. We also note that, for the incident pressures considered in this work, the results between ConWep and fluid-structure interaction modeling technique(Ganpule et al., 2013) are comparable (Supplementary Fig. S2).

Two types of boundary conditions were considered: (i) case-I: interaction of blast wave with full-body with regions below thorax fully constrained. This model mimics a typical warfighter with some kind of body shielding during occupational exposure (e.g., protection behind a wall, standing in a vehicle, breaching operation).(Kamimori et al., 2017; Wiri and Needham, 2016) This case also approximates the head surrogate studies using shock tubes. (ii) case-II: interaction of blast wave with full-body without any constraints. This model mimics a warfighter standing casually. To delineate the effects of blast wave transmission and global head motion (i.e., inertial effects) additional simulations were performed wherein the global head motion obtained from the above cases were used as an input to investigate the translation only and rotation only effects.

### 2.3 Solution Scheme

Nonlinear, transient, dynamic simulations were performed using an explicit scheme and massively parallel processing solver in LS-Dyna.(LSTC, 2017) Time history plots of acceleration and pressure are either filtered using SAE180 filter (to reduce the noise) or filter used in the experiment (for validation studies).

## 3. Results

### 3.1 Head Model Validation against Blast Loading

We validate the human head model against blast loading. We also compare the model response against experimentally measured *in vivo* strains in human volunteers. The details of validation are provided in the Supplementary Information.

### 3.2 Response of Full-Body and Head Subjected to Blast

Figure 2 shows the spatiotemporal evolution of kinematic response of the full-body model to the frontal blast of intensity 210 kPa for case-I (lower body constrained). Head undergoes translation in the anterior-posterior direction (i.e., the direction of blast) until ∼90 ms and retraces back to its original position by ∼200 ms (Fig. 2B-C). The linear displacement, measured at the center of mass, in the superior-inferior direction is small (peak displacement of 12 mm) as compared to the displacement in the anterior-posterior direction (peak displacement of 72 mm). In addition to the linear motion, the head undergoes rotation in the sagittal plane. Initially, the head undergoes flexion (i.e., it rotates clockwise about its center of mass). Peak flexion of 5°occurs at 15 ms (Fig. 2C) after which the head undergoes extension, peak extension of 24° occurs at 58 ms. After 60 ms, the head starts to rotate clockwise and retrace to the original position at 200 ms. The significant global head motion is not observed as the blast wave transcends the head and only noticeable after ∼3 ms. The acceleration profiles show a sudden increase followed by sharp decay (Fig. 2D), in response to the blast.

**Figure 2:**
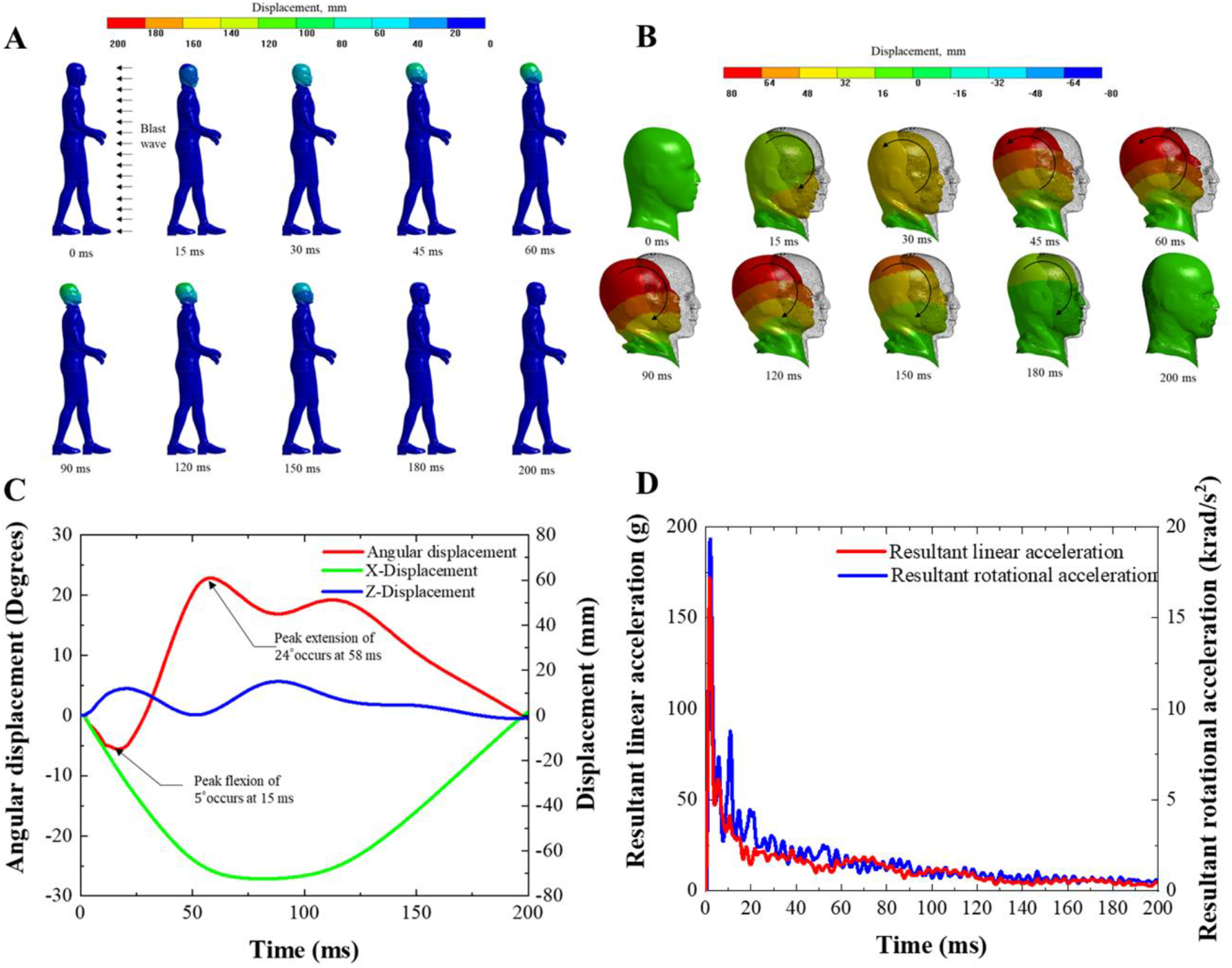
Spatiotemporal evolution of kinematic response of the human model to frontal blast for case-I (lower body constrained): Full-body (A), only head (B). Flexion (clockwise) and extension (anticlockwise) modes are indicated with arrows. Time history plots of linear, angular displacement (C), and head acceleration (D). Incident blast parameters correspond to peak overpressure of 210 kPa and positive phase duration of 6 ms.

For case-II (no lower body constraints), Fig. 3, linear motion is qualitatively similar to case-I, the magnitude of linear displacement is increased. The rotational motion is predominantly flexion until ∼60 ms, after which the head starts to retrace back towards its original position. The linear, angular displacements and angular velocity profiles are identical to case-I until ∼20 ms and deviate afterward due to the lack of extension of the head in case-II (Supplementary Fig. S8A, B). Peak head accelerations are identical in both cases (Fig. 2D, 3D) as peak head accelerations are established within the first few milliseconds. The peak head kinematics for all the simulated cases are tabulated in Table 1.

**Table 1:** Summary of peak biomechanical parameters for the head, brain, and lung for various blast intensities and boundary conditions considered in this work.

| Case I: Lower body constrained |  |  |  |  |  |  |
| --- | --- | --- | --- | --- | --- | --- |
| Incident overpressure, kPa | Peak resultant linear acceleration, $g$ | Peak resultant rotational velocity, rad/sec | Peak resultant rotational acceleration, krad/sec<sup>2</sup> | Peak coup pressure, kPa | Peak contre-coup pressure, kPa | Lung energy, J |
| 70 | 42.9 | 6.4 | 3.7 | 58 | -9.3 | 0 |
| 140 | 101 | 12.6 | 10.7 | 125 | -22.4 | 0 |
| 210 | 172 | 19.8 | 19.3 | 189 | -37.7 | 0 |
| Case II: No lower body constraints |  |  |  |  |  |  |
| Incident overpressure, kPa | Peak resultant linear acceleration, $g$ | Peak resultant rotational velocity, rad/sec | Peak resultant rotational acceleration, krad/sec<sup>2</sup> | Peak coup pressure, kPa | Peak contre-coup pressure, kPa | Lung energy, J |
| 70 | 42.0 | 4.8 | 3.7 | 58 | -9.3 | 2 |
| 140 | 102 | 9.2 | 10.6 | 125 | -22.4 | 18 |
| 210 | 172 | 14.3 | 19.3 | 189 | -37.7 | 45 |

**Figure 3:**
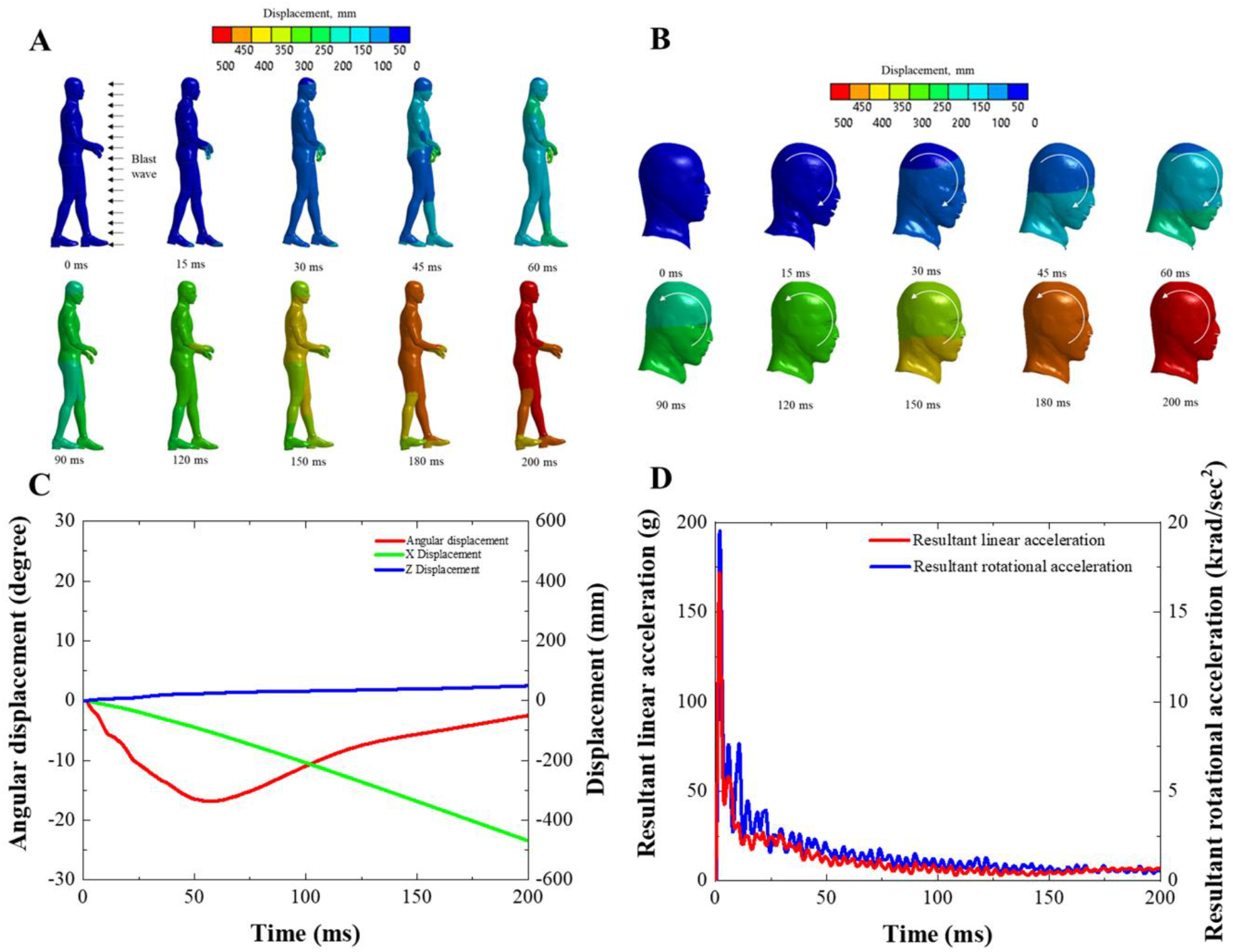
Spatiotemporal evolution of kinematic response of the human model to frontal blast for case-II (no lower body constraints): Full-body (A), only head (B). Flexion (clockwise) and retraction (anticlockwise) modes are indicated with arrows. Time history plots of linear, angular displacement (C), and head acceleration (D).

As the blast wave interacts with the head, the pressure is amplified due to the conversion of kinetic energy contained in a blast to the potential energy and part of the wave is transmitted to the brain. Figure 4 shows the spatiotemporal evolution of pressure (volumetric) and equivalent stress (deviatoric) in the brain for case-I, for the aforementioned loading. Pressure shows a typical coup-contrecoup pattern (Fig. 4A, B). Pressure wave dynamics in the brain play out within a few milliseconds (∼2 ms).

**Figure 4:**
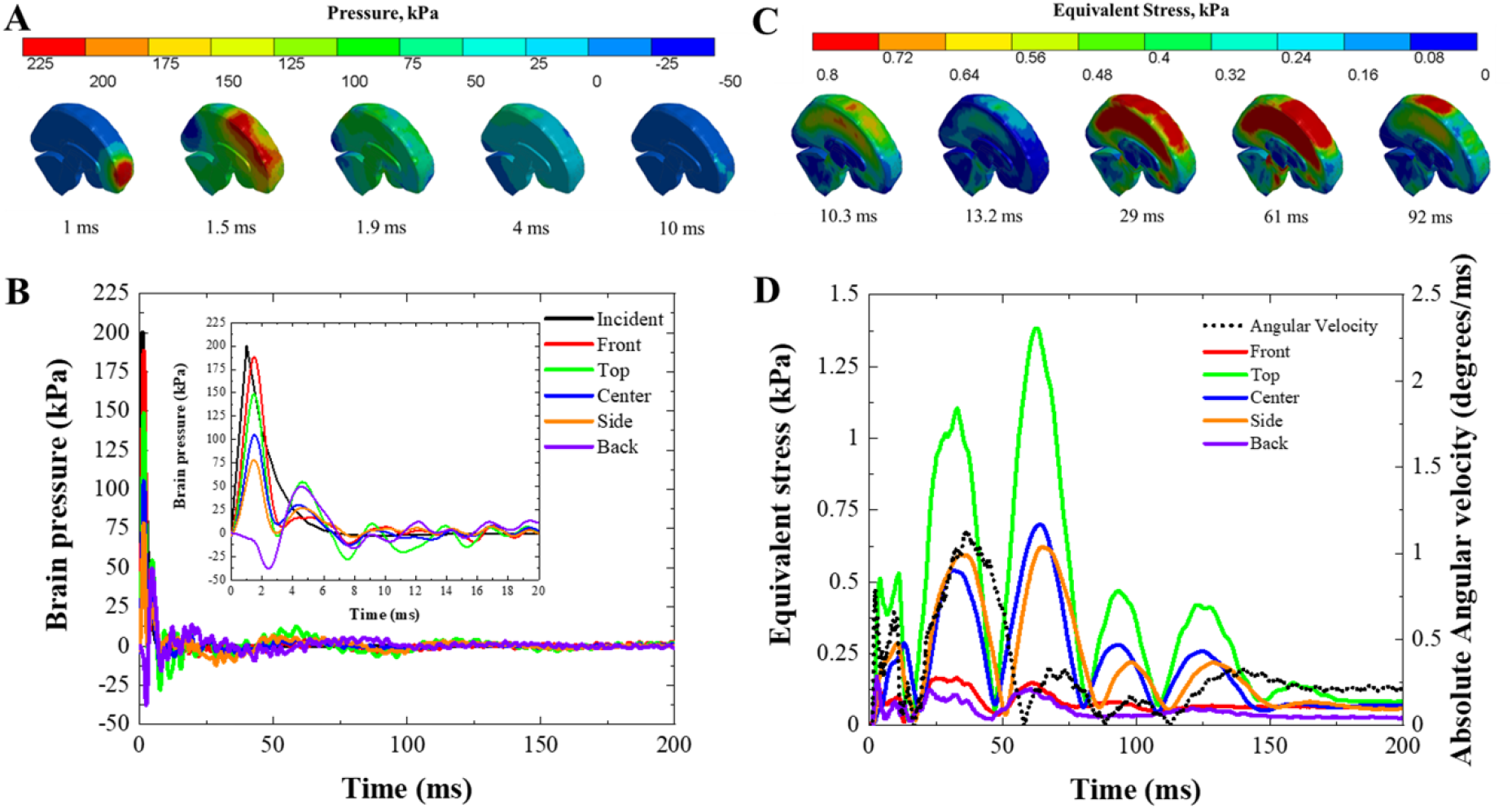
Kinetic response of the human model to frontal blast for case-I (lower body constrained). Spatiotemporal evolution of brain pressure (A, B) and equivalent stress (C, D). Pressure response shows a typical coup-contrecoup pattern; peaks are established within 1.5 ms. Equivalent stress shows a spherically converging pattern. Initial peaks are established within 10 ms; the pattern repeats itself (e.g., 10.3, 29, and 61 ms) corresponding to every head rotation and commensurate with the temporal evolution of angular velocity as depicted in C and D.

Equivalent stress evolution shows the spherically converging pattern that reaches the center ∼10 ms (Fig. 4C, D). Further, the temporal evolution of equivalent stress closely follows the temporal evolution of angular velocity, in terms of frequency of oscillation (Fig. 4D). This has two consequences: first, there is time-scale separation between volumetric and deviatoric response, and second, volumetric and deviatoric modes are governed by two different mechanisms. For case-II, pressure response (Supplementary Fig. S8C) is similar to case-I. Equivalent stress response (Supplementary Fig. S8D) is similar until ∼20 ms, after which response deviates considerably due to the lack of extension mode.

Results are qualitatively similar for other blast intensities (70 kPa, 140 kPa), peak values of key biomechanical parameters are tabulated in Table 1. The peak kinematic and pressure responses between case-I and case-II are almost identical, except for the angular velocity. The difference in angular velocity between these two cases is due to the presence of extension mode in case-I.

### 3.3 Contribution of Blast Wave Transmission and Global Head Motion to Stress Field

To delineate the effects of stress wave propagation and global head motion, we modified skull dipole (Fig. 1B) from deformable to rigid and prescribed computed displacements (from case-I) at the center of mass of rigid skull dipole. The locations of center of mass of full head and skull dipole are within 1%. Figure 5 shows the stress field in the brain for these cases. The wave propagation and translation of the head govern the volumetric response (Fig. 5A, B). Pure rotation alone governs the deviatoric response (Fig. 5C, D).

**Figure 5:**
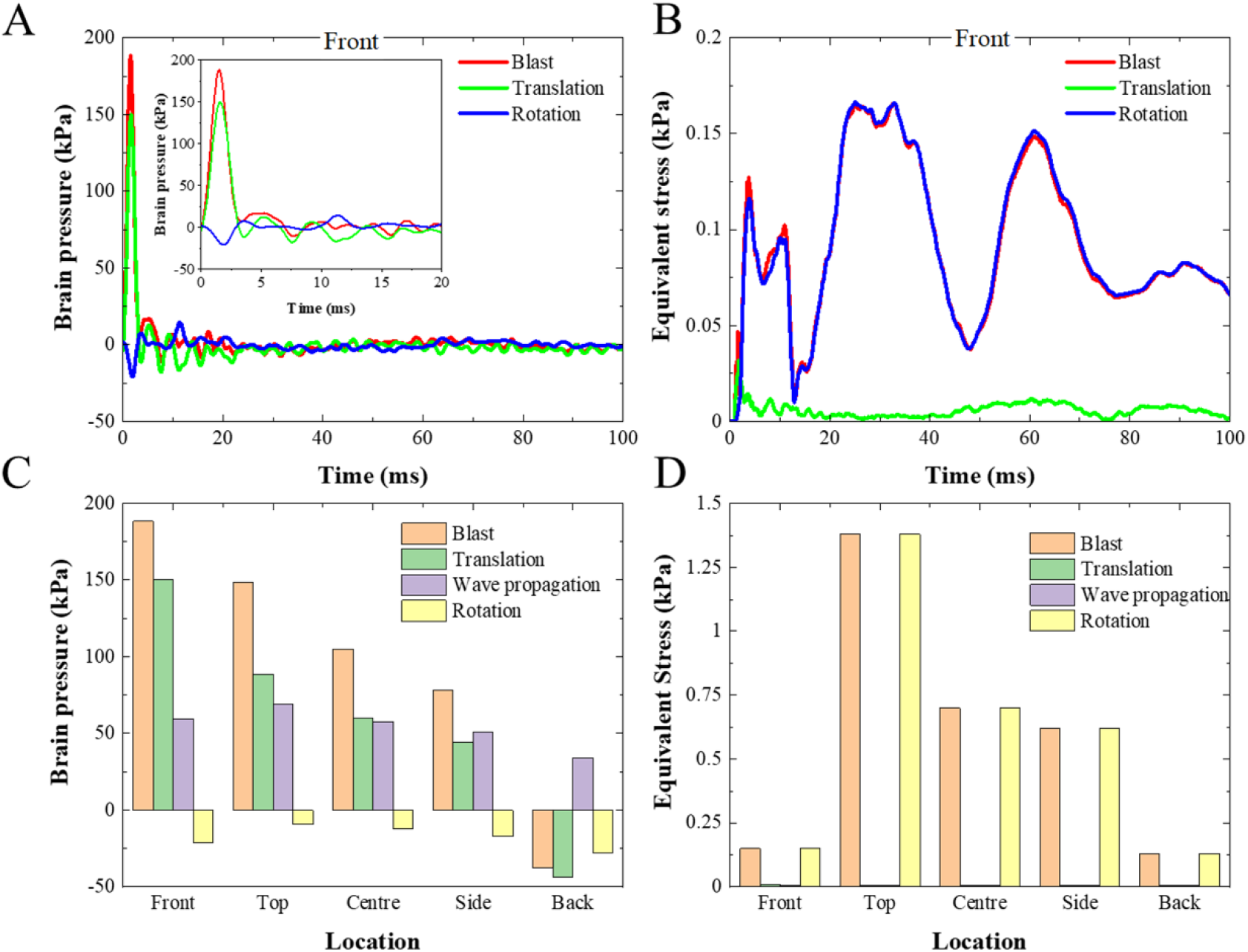
Estimates of contributions of wave propagation, translation, and rotation of the head to brain pressure and equivalent stress. Sample brain pressure-time (A) and equivalent stress-time (B) history plot at a front location in the brain. Peak brain pressures (C) and peak equivalent stress (D) for five locations are depicted in Fig. 1E. The contribution of wave propagation is obtained by subtracting the contribution of ‘translation and rotational only’ results from ‘full blast’ results.

The sensitivity of results to the material properties of the brain (Supplementary Fig. S9) shows similar overall trends as baseline response, in terms of the relative contribution of wave propagation, head translation, and head rotation.

### 3.4 Injured White Matter Fiber Tracts

Fig. 6 depicts the injured (damaged) white matter fiber tracts based on the axonal injury criterion of 13% tensile strain.(Giordano and Kleiven, 2014) White matter damage corresponding to blast and blast induced head rotation is almost identical with negligible damage in translation only case. Interestingly, the damage is diffuse.

**Figure 6:**
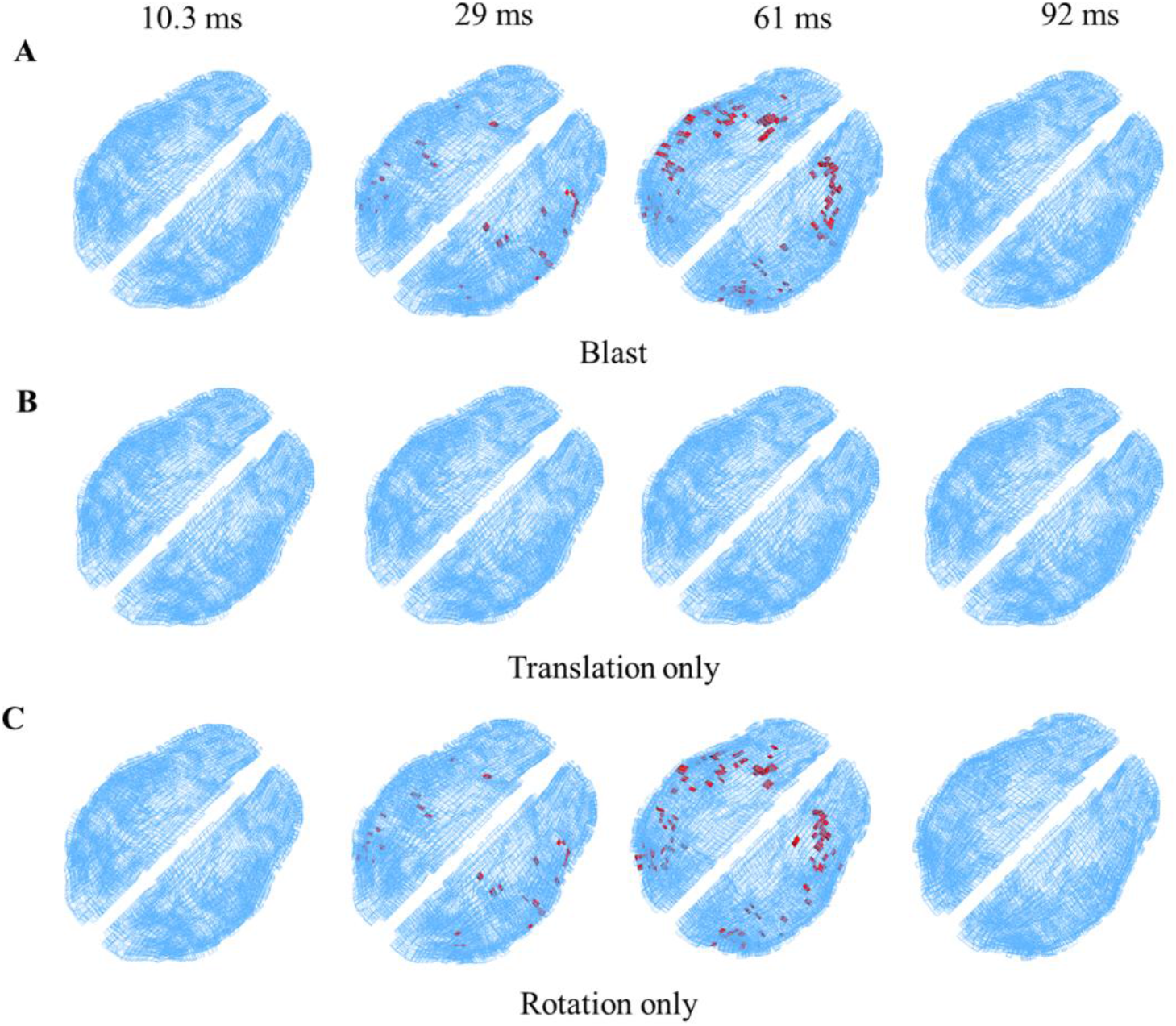
Injured regions (marked in red) of white matter fiber tracts exceeding axonal strain injury criterion: Full blast (A), only translation (B), only rotation (C). The damage is diffuse that encompasses regions of the corpus callosum, corona radiata, corticospinal tract, and internal capsule.

### 3.5 Strain Energy in the Lung

To explore the possibility of thoracic mechanism in transferring pressure pulse to the brain, we monitored strain energy in the lung (Table 1). Earlier investigations(Suneson et al., 1988; Suneson et al., 1990a, b) in this regard report that ∼700 J of work is required by the thoracic tissue to generate pressures of ∼125 kPa in the brain. The strain energy in the lung, corresponding to case-II, is 2, 18, 45 J, for blast intensity of 70, 140, and 210 kPa, respectively.

## 4. Discussion

We have simulated the response of a human against a primary blast by subjecting the model to static overpressure of three blast intensities (Supplementary Fig. S1) and investigated the resulting biomechanical response for 200 ms. Our results underscore a cascade of biomechanical response (Figs. 2-4). The early time response (<3 ms) is dominated by blast wave transmission and subsequent stress wave propagation within the skull and the brain. The response at a later time (5-200 ms) is dominated by global head motion. These results highlight the time-scale separation between the blast wave transmission effects and inertial effects and their direct correlation to generating mechanical fields within the brain. The wave propagation and linear acceleration of the head govern the volumetric response, whereas the rotational motion of the head governs the deviatoric response (Fig. 5). Further, the temporal evolution of deviatoric response within the brain is commensurate with the evolution of angular velocity (Fig. 4D). To the best of authors knowledge, we are the first to delineate these responses. This finding suggests that the aforementioned loading paradigms should trigger distinct and unique biochemical, pathological, and behavioural deficits.

Goldstein el al.(Goldstein et al., 2012) subjected the mice to blast loading. They found that mice underwent significant head rotation in response to blast and that resulted in neurodegeneration leading to learning and memory deficits; notably, learning and memory deficits were absent when the head was immobilized. Similar findings in rodents were reported by Risling et al.(Risling et al., 2011) and Aravind et al.(Aravind et al., 2020). Gullotti et al.(Gullotti et al., 2014) studied the response of the murine model to the blast and found that righting time deficit was correlated with the blast induced head acceleration, minimizing head acceleration eliminated righting time deficit. Thus, in addition to measurement of blast profiles, it is important to characterize head motions during blast exposure, to faithfully delineate effects of wave propagation, head translation, and head rotation.

Interestingly, in response to the blast, the head undergoes initial flexion, followed by an extension. For case-I (i.e., lower body constrained), we observe a peak flexion of 5° in 15 ms, followed by a peak extension of 24° at 58 ms (Fig. 2). For case-II (i.e., no lower body constraints), peak flexion of 17° is observed at 55 ms and significant extension is absent (Fig. 3). The flexion is seen because the center of mass of the whole body is at the hip, causing forward rotation. Further, the blast induces a larger thrust on the lower body than the head. These results are consistent, qualitatively, with experimental observation. Iwaskiw et al.(Iwaskiw et al., 2018) studied the response of PMHS filled with the brain simulant to a frontal blast and observed initial flexion (peak flexion of ∼3°) followed by an extension (peak extension of ∼45°). The peak magnitudes from this work are not directly comparable to Iwaskiw et al.,(Iwaskiw et al., 2018) as experimental conditions are different (and may be affected by shock tube end effects) from the current simulations. The flexion and extension modes cannot be deduced from the validation experiments used for model validation.(Levine et al., 2014; Skotak et al., 2018). Note that simulated cases (i.e., case-I, II) represent two extremes and battlefield primary blast scenarios are bounded within these extremes.

Our results suggest that out of various proposed mechanisms, wave transmission and head rotation are dominant mechanisms. The qualitative (Fig. 2-4) and quantitative (Table 1) responses of the head and brain are comparable to responses in the blunt impact biomechanics literature.(Giordano and Kleiven, 2014; Hajiaghamemar et al., 2020; Hernandez and Camarillo, 2019; Hernandez et al., 2015a; Hernandez et al., 2015b; Ji et al., 2015; Ji et al., 2014; Mao et al., 2013; Patton et al., 2012; Rowson et al., 2012; Zhang et al., 2004) This suggests that the blast induces a similar biomechanical cascade (e.g., coup-contrecoup pressure and spherically converging shear wave patterns) and peak responses as that of blunt impact. However, the duration of acceleration and pressure profiles are typically smaller (<5 ms in blast, 10-20 ms in impact).

Peak negative (contrecoup) pressures for all intensities (Table 1) are lower than the widely postulated cavitation threshold of −100 kPa.(Panzer et al., 2012) At higher intensities, the possibility of negative pressure going below −100 kPa in our model cannot be precluded. Relating the value of negative pressure to the onset of cavitation in the brain tissue is nontrivial.(Barney et al., 2020)

The computed strain energy in the lung (2-45 J), corresponding to case-II, is considerably lower than the energy required by the thoracic tissue (∼700 J) to generate significant pressures in the brain (∼125 kPa), due to the thoracic compression. Yet, at these energies, the possibility of generating vascular pathology due to the inflation of veins and arteries(Dagro and Ramesh, 2019; Rubio et al., 2020) cannot be ruled out. The macroscopic model used in this work is insufficient to investigate the effects of the vasculature.

The sensitivity of brain response (Supplementary Fig. S9) to high strain rate material properties implied in the blast is restricted to generating more oscillatory response (corresponding to higher frequencies) and faster relaxation of stress in the brain without altering the basic biomechanical cascade. The role of high strain rate response and associated fluctuations in the mechanical field of the brain in causing injury(Abdul-Wahab et al., 2020; Bar-Kochba et al., 2016; LaPlaca et al., 2005; Skotak et al., 2013) is not well understood. Our results underscore that work in this regard will be valuable to fully understand the bTBI.

The blast induced head rotation alone produces diffuse injury pattern in the white matter fiber tracts. We predict significant white matter damage for blast intensity of 210 kPa, minor damage for the intensity of 140 kPa, and no damage for the intensity of 70 kPa. The injured regions, corresponding to blast intensity of 210 kPa, are diffuse in nature encompassing regions of the corpus callosum, midbrain, corticospinal tract, corona radiata, and internal capsule. Thus, the blast also produces diffuse injury pattern encompassing regions that are reported in blunt impact events (e.g., Ganpule et al.(Ganpule et al., 2017)). The most commonly reported neurological abnormalities (e.g., Phipps et al.(Phipps et al., 2020) and references therein) in the bTBI population are white matter irregularities and diffuse axonal injury. Clinical investigations comparing cohorts with a history of blast and blunt impact TBI are limited. Dennis et al.(Dennis et al., 2018) and Dretsch et al.(Dretsch et al., 2015) respectively carried out neuroimaging and neuropsychological studies in deployed U.S. Army soldiers specifically identified for blast and blunt impact injuries. They found that there were no statistically significant differences between the blast and blunt impact injury participants. These findings reinforce the role of blast induced head rotation in bTBI cohorts.

In this study, the detailed full human body model of GHBMC is implemented to investigate the direct cranial transmission and global head motion to field-relevant blast loads. Still, the bio-fidelity of current GHBMC model can be further improved by incorporating region-specific skull and facial bones, the details of sulci, gyri and brain vasculature, modifying the linear viscoelastic brain material to non-linear viscoelastic and improving the mesh resolution of head components. Strain measurements in PMHS or human under blast loading are not available. Experimental strain measurements under impact loading are only available for mild, non-injurious acceleration in human volunteers. Hence, the model cannot be validated under injurious, experimental measurements of strain. Additionally, despite the neck model was validated for local tissue kinetics and global head kinematic response under quasi-static (flexion, extension, axial rotation) and/or dynamic (frontal, lateral, rear impact) loading scenarios, the predicted rotational acceleration response to blast loads was less definitive (Supplementary Fig. S3). Thus, on one side, there is scope to further improve the head and neck bio-fidelity in the existing model. On the other side, there is an opportunity for experimentalist to conduct the measurements of rotational head accelerations under blast loads which will be great source for head model validation. Note that, the neck is the most complex region to model and existing neck models are evolving. Despite the forementioned improvement areas and limitations of head-neck models, the findings reported in this study are not affected by them.

## 5. Conclusion

The response of human under the primary blast loading is studied using a detailed, full-body computational model. Results elucidate biomechanical cascade of events in human. The key findings are:

1. The response of the brain to the blast is governed by blast wave transmission effects and inertial effects that generate distinct biomechanical fields within the brain. Blast transmission and linear motion of the head govern the volumetric response, whereas the rotational motion of the head governs the deviatoric response.
2. The boundary conditions of the lower body (i. e., constrained vs. free) play an important role in governing temporal evolution of deviatoric response within the brain due to the additional rotational mode (i.e., head extension) in constrained lower body case. Volumetric response is unaffected by the boundary condition as the volumetric response is played out within very short time (<5 ms) for which the head rotational modes are negligible.
3. There are more commonalities to the biomechanical response of the brain under blast and blunt impact loading scenarios than the differences. Our results demonstrate the similarities in terms of coup-contrecoup pressure and spherically converging shear wave patterns and their magnitudes, observed peak accelerations, and diffuse nature of axonal damage.
4. Blast induced head rotation alone produces a diffuse injury pattern in white matter fiber tracts.

Overall, these results provide unique mechanistic insights into brain biomechanics under the primary blast. These findings have great utility in designing of next-generation personal protective equipment and will augment ongoing bTBI research using animal models and clinical studies.

## Acknowledgments

Authors acknowledge Global Human Body Model Consortium-owned GHBMC Model distributed to IIT Roorkee through a license agreement with Elemance.

