## Supplementary Material for "In silico investigation of biomechanical response of a human subjected to primary blast"

### Supplementary Information for "In silico investigation of biomechanical response of a human subjected to primary blast."

#### Supplementary text:

#### Full-body Human Model

Human model (Fig. 1) was created from a 50$^{th}$ percentile healthy male volunteer (age = 26 years, weight = 78.6 kg, height = 174.9 cm) using MRI and CT imaging.(Gayzik et al., 2011b, 2011a, 2009) The subject matched within 5% of fifteen anthropometric targets from a survey by the U.S. Army. The resulting CAD dataset included 418 parts containing 179 individual bones, 46 organs, 96 muscles, 37 blood vessels, and 26 ligaments, tendons, and cartilaginous structures. CAD dataset was divided into five body region models head, neck, thorax, abdomen, and pelvic-lower extremity. The finite element model for each of five body regions (Fig. 1A) has been developed separately, through multi-institutional effort, using high quality mesh (mostly hexahedral). Body region models were integrated into the full-body model using appropriate head-neck, neck-thorax, thorax-abdomen, abdomen-pelvic interfaces (either shared nodes or no slip, no penetration contact). CAD and FE models were developed with the objective of incorporating macroscopic (continuum) level features. The full-body model contains ~1.25 million nodes and ~2.5 million elements.

#### Head Model

The head model,(Mao et al., 2013) Fig. 1B, consisted of the scalp, skull, facial bones, sinuses, dura, pia, arachnoid, falx, tentorium, bridging veins, and brain. The brain was segmented into the cerebrum, cerebellum, ventricles, brainstem, thalamus, corpus callosum, and basal ganglia. The cerebrum was further segmented into white and gray matter. The model was meshed using 270,552 high quality elements. The skull and facial bones were modeled as elastic-plastic, whereas the brain and scalp were modeled as linear, viscoelastic. The rest of the components were modeled as linear, elastic, consistent with the literature.(Ganpule et al., 2017; Giordano and Kleiven, 2014; Ji et al., 2015) For baseline simulations, we kept all the material properties (including brain, supplementary Table S2) same as that of the original model,(Mao et al., 2013) as the model was validated under these set of properties. In addition to baseline simulations, we have incorporated the hyperviscoelastic constitutive model and associated material parameters for the brain tissue that are valid over the strain rate regime relevant for the blast (up to $10^6$ $s^{-1}$); the details can be found in Chafi et al.,(Chafi et al., 2011) the material properties are listed in Supplementary Table S3. We note that results using Chafi et al.(Chafi et al., 2011) brain material properties only appear in supplementary material and provide a few additional insights.

**Neck Model**

Neck model (Fice et al., 2011; Panzer et al., 2011) consisted of cervical spine, active and passive muscle elements, skin, and adipose tissue. The cervical spine comprised the cervical vertebrae (C1–C7) with the associated cortical and cancellous bones, ligaments, intervertebral discs, endplates, nucleus pulposus and facet joint cartilages. The relative position of the tissues in the neck model was determined from the CT and MRI data. The ligamentous cervical spine was integrated with 25 pairs of 3D passive and 1D active muscles, adipose tissue, and skin. The neck model was appropriately meshed using solid, shell, and beam elements. The various tissues of the neck were represented using appropriate material models. Material properties were obtained from experiments or adopted from the literature. Some of the tissues of the neck also contained failure models such as hard tissue fracture, progressive failure for ligament, and intervertebral disc avulsion. The neck model was validated for local tissue kinetics and global head kinematic response under quasi-static (flexion, extension, axial rotation) and/or dynamic (frontal, lateral, rear impact) loading scenarios (see Supplementary Table S1). Experimental data in human volunteers and PMHS were used for whole neck validation.

**Incorporation of White Matter Structural Anisotropy into Head Model**

White matter structural anisotropy in the form of fiber orientations was incorporated (Fig. 1C) using diffusion tensor imaging (DTI). DTI data is based on the DTI of a 25-year-old healthy male volunteer.(Ganpule et al., 2017) DTI data were aligned (Fig. 1D) with the finite element data using appropriate rigid body translation and rotation. The skull and cerebral white matter of DTI dataset aligned perfectly with respective components of the head model and hence no scaling was required. The resolution of DTI data was 2 $mm^3$, whereas the resolution of finite element mesh was 2-3 $mm^3$. To compensate for the slight difference in the resolution, the orientations of DTI data points contained within an element were averaged. For simplicity, we have only considered structural anisotropy of white matter and material response is modeled as an isotropic. Strain along fiber direction was computed from Green-Lagrange strain tensor ($E$) during postprocessing, using equation 1. This approach is reasonable and justified in this work, as the main focus is to investigate the biomechanical response under the primary blast. Zhao and Ji(Zhao and Ji, 2018) found that computed axonal strains were statistically similar when fiber dispersion for individual fibers was explicitly considered using an anisotropic constitutive model and when it was set to 1/3, representing isotropy. In another study, Ji et al.(Ji et al., 2015) have demonstrated that computing axonal strains during postprocessing provides reasonable qualitative agreement with the DTI based clinical findings of sports concussion.

$$\varepsilon_{fiber} = \bar{l}_{el}^{\;T} \cdot E \cdot \bar{l}_{el} \quad [1]$$

where, $\bar{l}_{el}$ is a vector representing fiber orientation and $E$ is Green-Lagrange strain tensor.

#### Head Model Validation against Blast Loading

In this work, we validate the human head model for blast loading. Skotak et al.(Skotak et al., 2018) the response of facial and ocular countermeasures safety headform mounted on Hybrid-III neck to frontal blast. They reported peak headform acceleration (measured at the center of mass) and reflected pressures at different locations on the surface of the headform along the mid-sagittal plane. In another study, Levine et al.(Levine et al., 2014) measured linear and rotational acceleration of Hybrid-III head during frontal blast exposure of Hybrid-III mannequin (head, neck, and thorax) in standing position. Bir(Bir, 2011) studied the response of fresh, post-mortem human subjects (PMHS) to frontal blast; reflected pressures on the surface of the head and intracranial pressures in the brain were reported for incident overpressures of ~70 kPa and ~100 kPa. We simulated these experiments and compared the model response against measured biomechanical data (Supplementary Fig. S3-S5, Supplementary Table S4-S5). The obtained kinematic (i.e., head acceleration) and kinetic (i.e., reflected and intracranial pressures) response from the simulations match reasonably well with the experiments and commensurate with current acceptable standards in head injury biomechanics literature.(Garimella et al., 2018; Miller et al., 2017)

**Model Validation against Brain Strains:**

Knutsen et al.,(Knutsen et al., 2020) using tagged MRI, measured the full-field brain strains in human volunteers during mild neck rotation (peak head acceleration ~200 rad/s$^2$) and neck extension (peak head acceleration ~330 rad/s$^2$) using a custom made, MRI-compatible head motion apparatus. The apparatus fits inside the bore of an MRI machine and guides voluntary head motion. This results in axial and sagittal plane rotations of the head for neck rotation and neck extension, respectively. To simulate the experiment, the measured angular displacement-time history was used as an input to the simulations. Supplementary Fig. S6-S7 show the comparison of strains between a model and experiment. The agreement between a model and experiment is reasonable. The model is able to capture the peak strain magnitudes as well as distribution of strain.

**Supplementary Figures:**

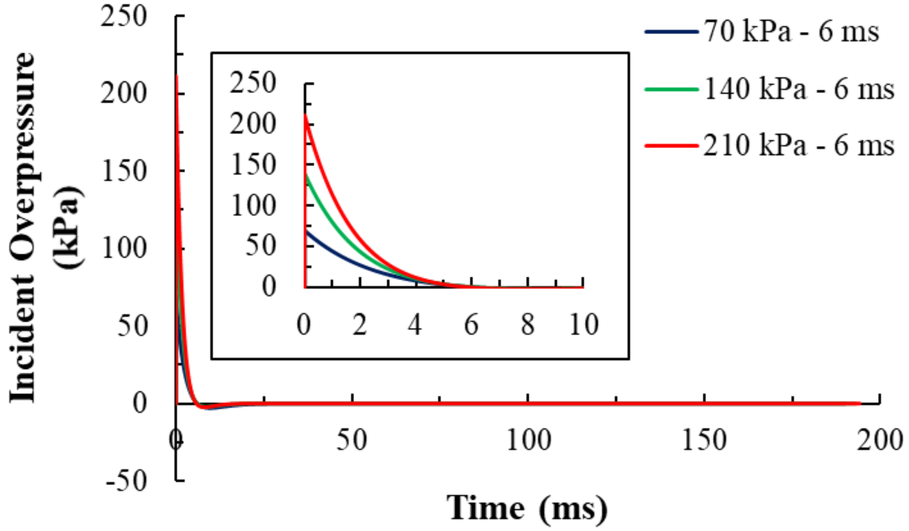


**Supplementary Fig. S1.** Incident (static) overpressure-time profiles of blast wave. We have considered three blast intensities with peak incident overpressures of ~70, ~140, ~210 kPa, and a positive phase duration of ~6 ms. Incident overpressure-time histories follow a typical Friedlander profile, with a sharp rise followed by nonlinear decay. There is no other input to the model other than the incident overpressure.

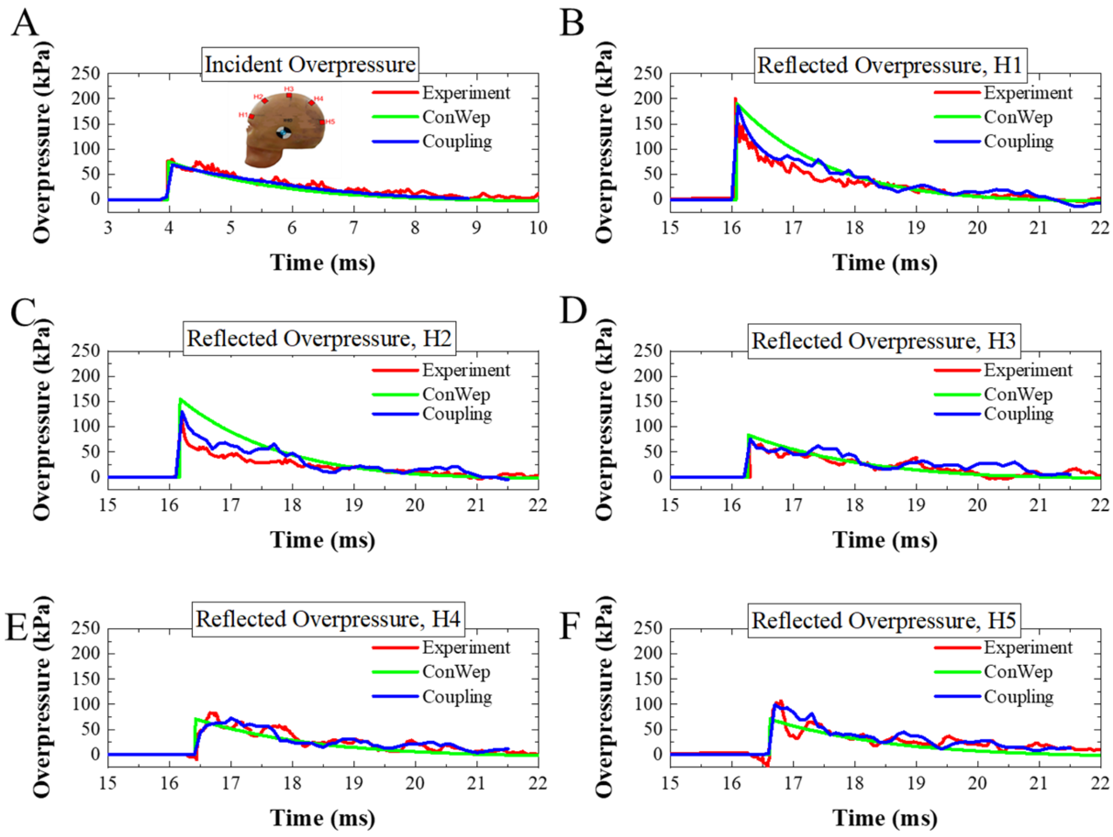


**Supplementary Fig. S2.** Comparison of ConWep and Coupling (fluid-structure-interaction) modeling techniques in simulating blast wave-head interactions. The pressure-time histories of the incident and reflected pressures at various locations on the head are shown. Insert of panel A shows the locations on the head where the reflected pressures are measured in the experiment.(Skotak et al., 2018) ConWep modeling technique is computationally efficient as it does not require explicit modeling of the air domain between the source and the target. It is especially attractive to simulate the interaction of the blast wave with the full-body human as the Coupling approach requires at least 10-fold computational cost and powerful computational facilities.

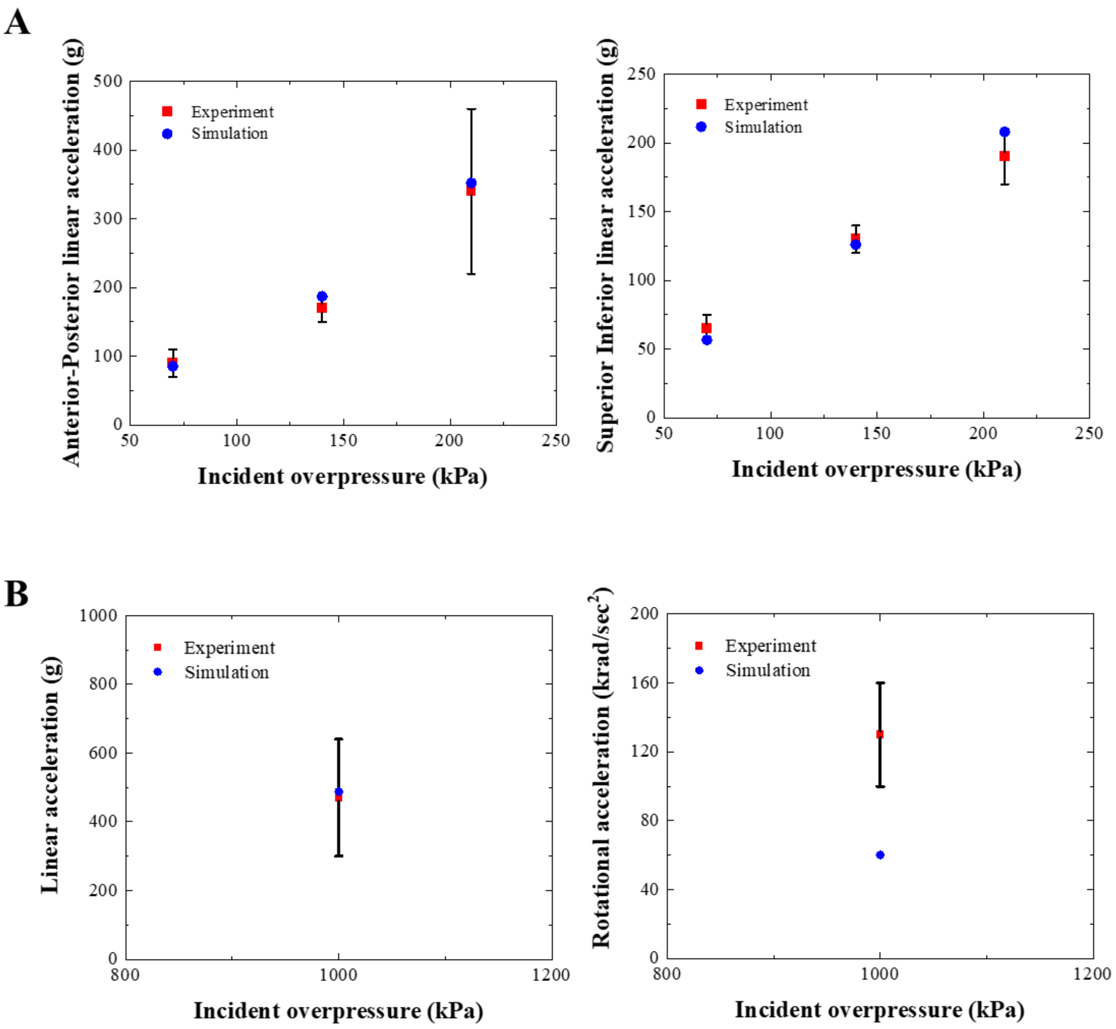


**Supplementary Fig. S3.** Comparison of simulated head acceleration against the experiments of Skotak et al.(Skotak et al., 2018) (A) and Levine et al.(Levine et al., 2014) (B). Skotak et al.(Skotak et al., 2018) conducted experiments on facial and ocular countermeasures safety headform using a shock tube with specimen placed in the shock tube test section. Their results from shock tube experiments are comparable to subsequent field trials,(Chandra et al., 2017) using actual explosives, by the same group. We have used shock tube results for validation as they are more comprehensive than the field trials. Using a blast chamber facility, Levine et al.(Levine et al., 2014) conducted the experiments on a Hybrid-III mannequin (head, neck, and thorax) in a standing position subjected to 1 kg of C4 kept at distances of 1 m. Note that the peak rotational accelerations from Levine et al.(Levine et al., 2014) are less definitive as sensors reached their design limit.

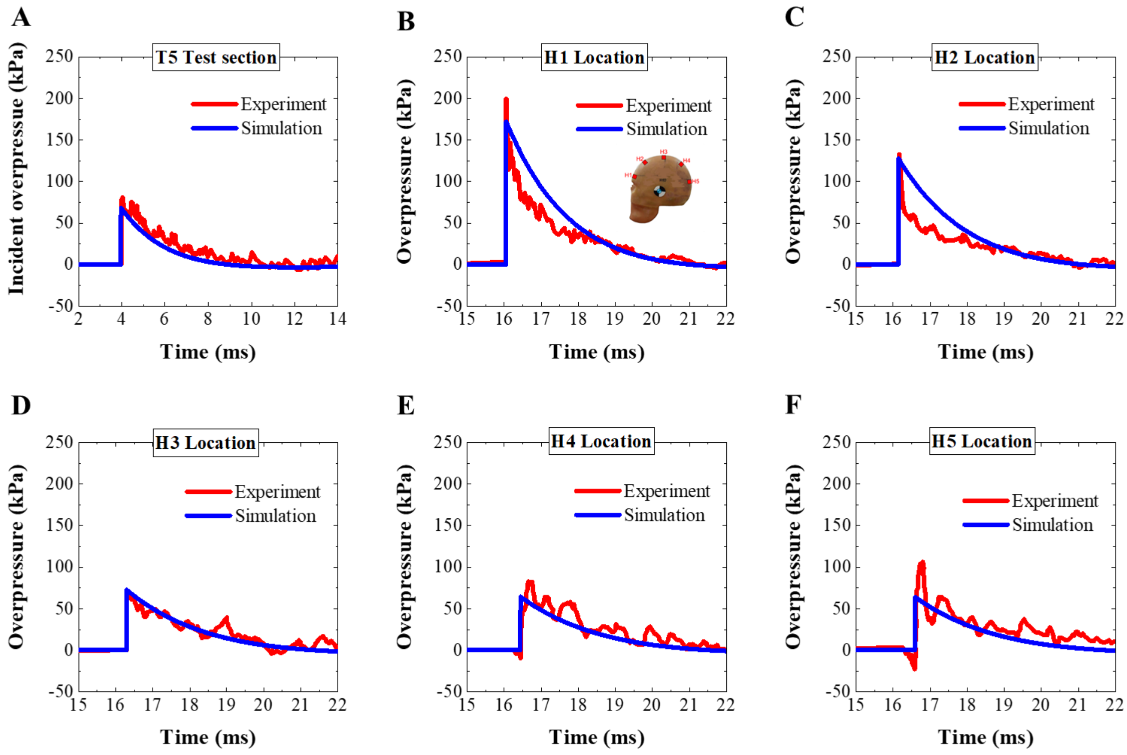


**Supplementary Fig. S4.** Comparison of the simulated incident (A) and reflected (B-F) pressures against the experiments of Skotak et al.(Skotak et al., 2018) Reflected pressure measurement locations are shown in the inset of panel 'H1 Location'. Corresponding correlation and analysis (CORA) scores are given in Supplementary Table S4.

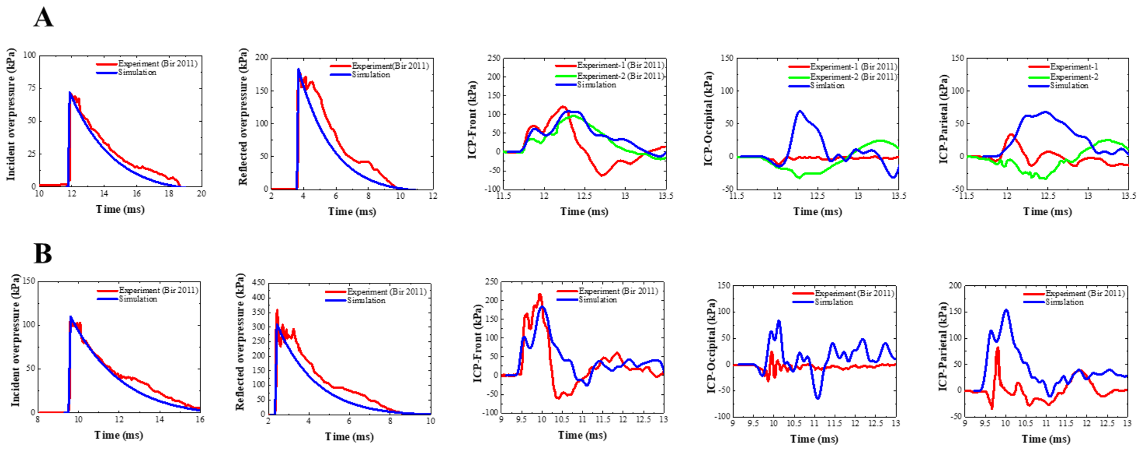


**Supplementary Fig. S5.** Comparison of the simulated incident, reflected, and intracranial (ICP) pressures against the experiments of Bir et al.(Bir, 2011) in PMHS conducted using a shock tube. CORA scores for ICP are given in Supplementary Table S5. Note that the CORA scores for ICP are small due to the inherent challenges associated with repeatable measurements of brain pressures (ICP profiles of Experiment-1 and Experiment-2 deviate considerably).

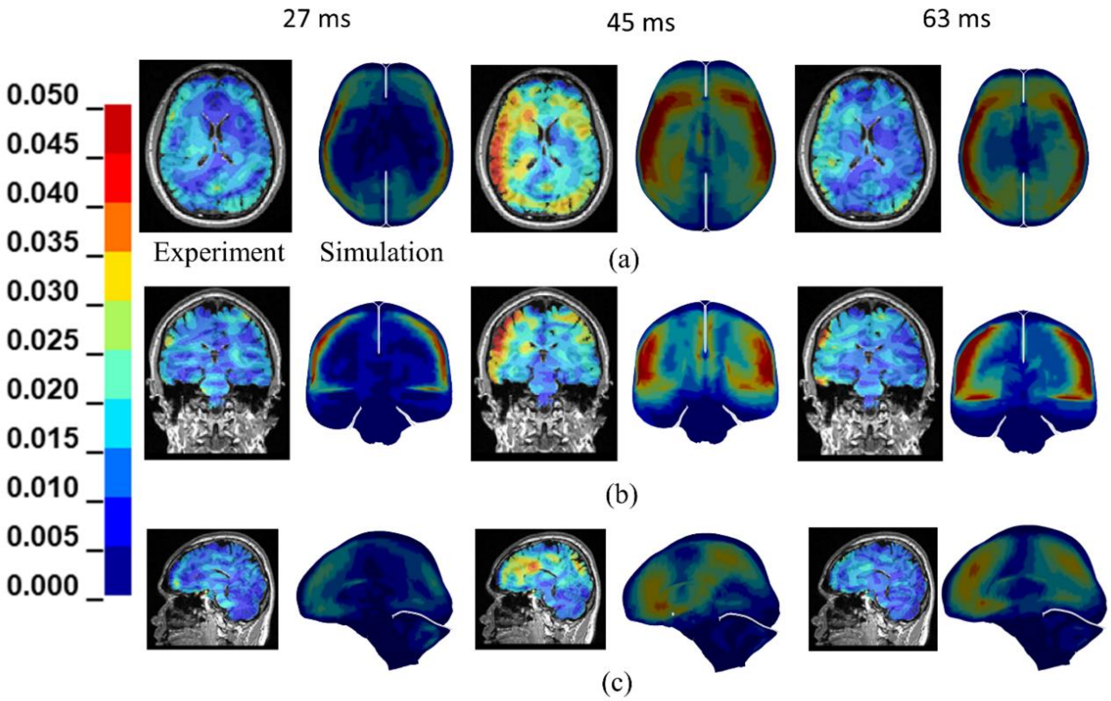


**Supplementary Fig. S6.** Comparison of maximum principal strain from the model with the head-neck rotation experiments of Knutsen et al.(Knutsen et al., 2020) (a) axonal, (b) coronal and (c) sagittal plane views. The agreement between a model and experiment is reasonable in terms of the peak strain magnitude and distribution of strain.

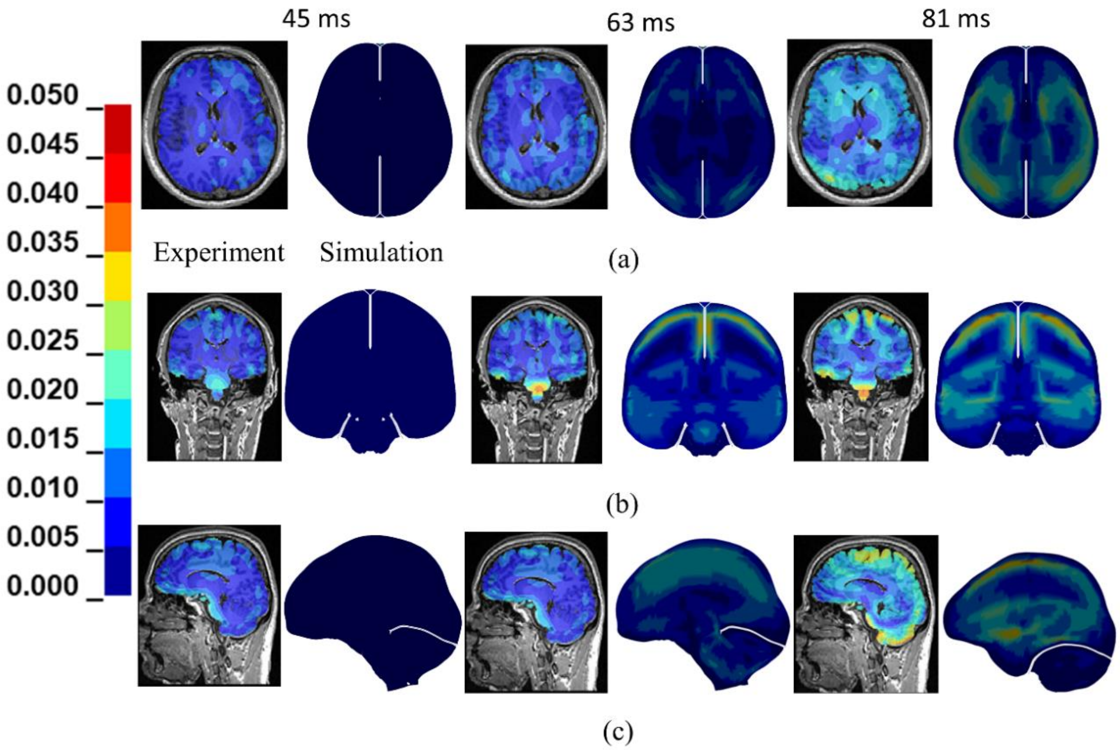


**Supplementary Fig. S7.** Comparison of maximum principal strain from the model with head-neck extension experiments of Knutsen et al.(Knutsen et al., 2020) (a) axonal, (b) coronal and (c) sagittal plane views. The agreement between a model and experiment is reasonable in terms of the peak strain magnitude and distribution of strain.

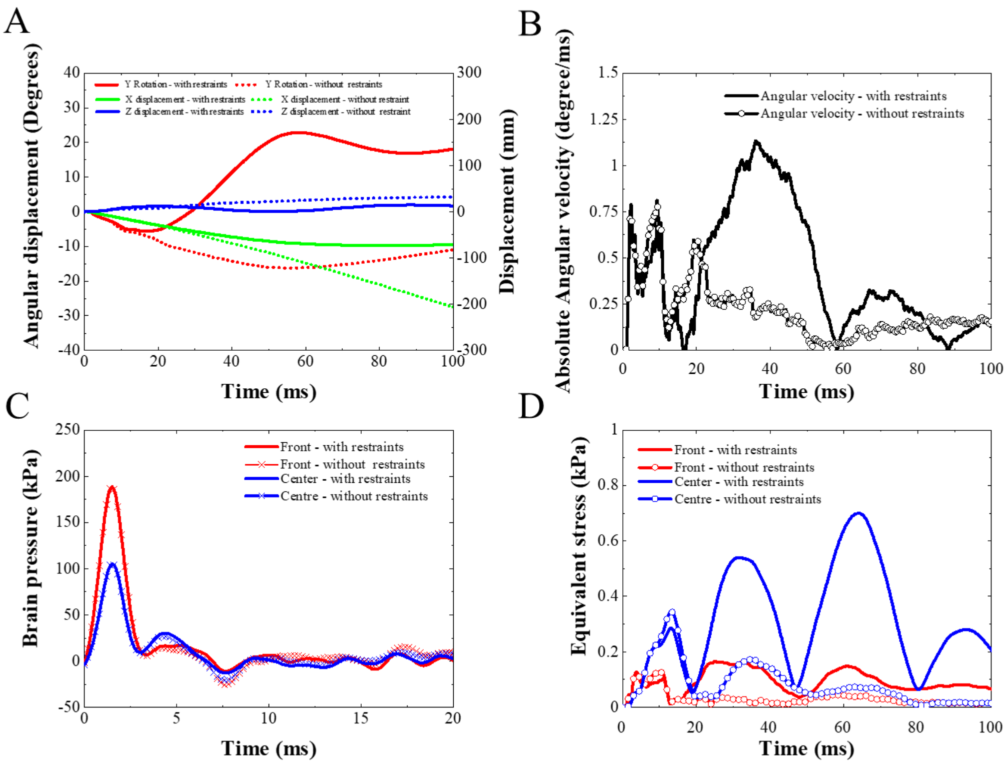


**Supplementary Fig. S8.** Comparison of kinematic and kinetic response for case-I (lower body constrained) and case-II (no lower body constraints): linear and angular displacement (A), angular velocity (B), brain pressure (C), equivalent stress (D).

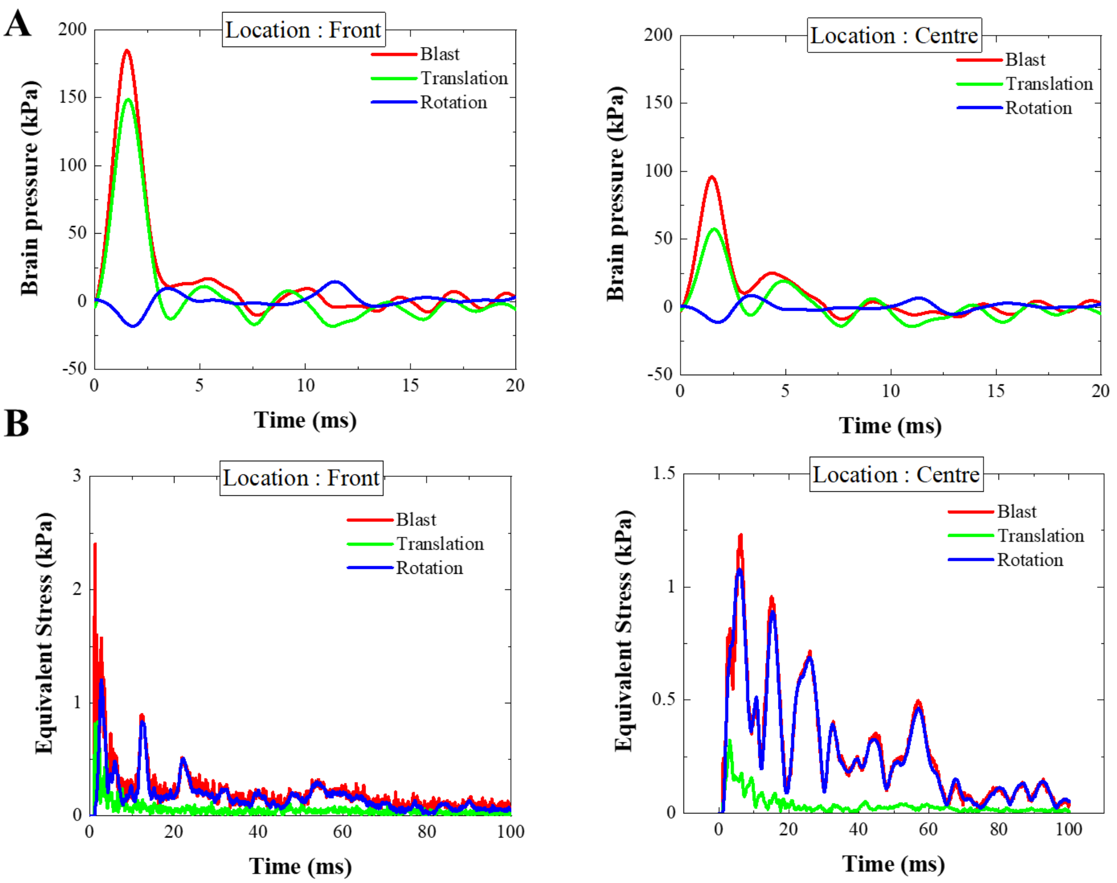


**Supplementary Fig. S9.** The sensitivity of results to the material properties of the brain show similar overall trends as baseline response. In this case, the material properties incorporate high strain rate response (see Supplementary Table S3). The volumetric response is comparable to the simulations with baseline properties (Fig. 5). Deviatoric response shows larger fluctuations corresponding to the high strain rate material properties. Considerable stress relaxation is also evident due to larger viscous dissipation and relaxation over wider time regimes.

#### Supplementary Tables:

**Supplementary Table S1.** Summary of validation of the full-body human model. Model is validated at full body and organ levels.

| Serial Number | Study | Full-body human model | Focus of validation | | | | | | Source of experimental data |
|---|---|---|---|---|---|---|---|---|---|
| | | | FBM | RBM | | | | | |
| | | | | Head | Neck | Thor | Abdo | PLX | |
| 1 | Mao et al.(Mao et al., 2013) | - | - | x | - | - | - | - | Brain response from frontal impact test<br>(Nahum et al. 1977)<br>Brain response from frontal impact test<br>(Trosseille et al. 1992)<br>Brain-Skull motion test<br>(Hardy et al. 2001, 2007)<br>Skull response<br>(Yoganandan et al. 1995, Hodgson et al. 1970)<br>Facial response from frontal impact test<br>(Nyquist et al. 1986, Allsop et al. 1988)<br>Brain contusion from frontal impact test<br>(Nahum et al. 1976) |
| 2 | Fice et al.(Fice et al., 2011) | - | - | - | x | - | - | - | Volunteer rare impact sled test<br>(Davidsson et al. 1998)<br>PHMS rare impact sled test (Deng 1999) |
| 3 | Panzer et al.(Panzer et al., 2011) | - | - | - | x | - | - | - | Flexion and extension response<br>(Nightingale et al. 2002, 2007; Wheeldon et al. 2006)<br>Ligament strain<br>(Punjabi et al. 2004, Ito et al. 2005, Bolton et al. 2006) |
| 4 | Poulard et al.(Poulard et al., 2015) | - | - | - | - | x | - | - | Rib segment anterior– posterior bending<br>(Kindig et al. 2009)<br>Point loading of the ribcage<br>(Kindig et al. 2013)<br>Frontal pendulum impact test<br>(Kroell and Schneider, 1971;<br>Lebarbe and Petit 2012)<br>Lateral impact test<br>(Shaw et al. 2006)<br>Table top test (Kent et al. 2004) |
| 5 | Li et al.(Li et al., 2010) | - | - | - | - | x | - | - | Rib fracture experiments<br>(Li et al. 2010) |
| 6 | Hayes et al.(Hayes et al., 2014) | - | - | - | - | x | - | - | Frontal sled tests<br>(Forman et al. 2006)<br>Lateral impact tests<br>(Pintar et al. 1997)<br>Lateral impact tests (Maltese et al. 2002) |
| 7 | Beillas and Berthet(Beillas and Berthet, 2012) | - | - | - | - | - | x | - | Rigid bar impact to abdomen<br>(Hardy et al. 2001) |
| 8 | Yue et al.(Yue and Untaroiu, 2014) | - | - | - | - | - | - | x | PMHS knee impact tests<br>(Balasubramanian et al. 2004)<br>human leg and thigh in dynamic latero-medial bending tests<br>(Kerrigan et al. 2004)<br>knee–thigh–hip response in frontal impact tests<br>(Rupp et al. 2003) |
| 9 | Untaroiu et al.(Untaroiu et al., 2013) | - | - | - | - | - | - | x | Femur compression test<br>(Keyak et al. 1998, 2001)<br>Femer three point bend test<br>(Funk et al. 2004)<br>Bending and compression test<br>(Ivarsson et al. 2008)<br>Bending and compression test<br>(Untaroiu et al. 2008) |

| | | | | | | | | | |
|---|---|---|---|---|---|---|---|---|---|
| 10 | Shin et al.(Shin and Untaroiu, 2013) | - | - | - | - | - | - | x | Axial impact, inversion/eversion tests (Funk et al. 2000, 2002) |
| 11 | Aira et al.(Aira et al., 2019) | M50-O | x | - | - | x | - | - | Costovertebral rib response Duprey et al. (2010); Intervertebral joint (Panjabi et al. (1976)); Rear pendulum impact (Forman et al. (2015); blunt lateral impact to upper extremity (Koh et al. 2005) |
| 12 | Schap et al.(Schap et al., 2019) | M50-O | x | x | - | x | - | x | Frontal sled PMHS tests (Shaw et al. 2009) |
| 13 | Gaewsky et al.(Gaewsky et al., 2019) | M50-O | | x | | x | | | Volunteer kinematic test data (Buhrman 1998) |
| 14 | Somasundaram et al.(Somasundaram et al., 2019) | M50-O | x | - | - | x | - | - | Spine response corridors from WSU UBB PMHS tests |
| 15 | Weaver et al.(Weaver et al., 2018) | M50-O | - | - | - | - | - | x | Pelvis impactor test (Bouquet et al. 1994,1998) |
| 16 | Devane et al. (Devane et al., 2019) | M50-O | x | x | x | x | x | x | Low speed frontal sled test volunteer, PMHS data (Beeman et al. 2012) |
| 17 | Zheng et al.(Zheng et al., 2018) | - | | - | - | x | - | - | Cadeveric tests on lumbar spine (Demetropoulos et al. 1998) |
| 18 | Decker et al.(Decker et al., 2017) | M50-O | x | x | - | - | - | - | Frontal/lateral impact human volunteer tests (Wismans et al. 1986) |
| 19 | Xiao et al.(Xiao et al., 2017) | M50-O | - | - | - | x | - | - | Frontal sled PMHS tests (Shaw et al. 2009) |
| 20 | Park et al.(Park et al., 2016) | M50-O | x | - | - | x | - | - | Lateral sled PMHS tests (Lessley et al. 2010) |
| 21 | Arun et al.(Arun et al., 2016) | M50-O | x | x | x | x | x | x | Far side lateral impact tests (Pintar et al. 2007) |
| 22 | Kindig et al.(Kindig et al., 2015) | M50-O | - | - | - | x | - | - | Quasi-static point loading on PMHS ribcage (Kindig et al. 2010) Dynamic loading on PMHS ribcage (Vezin and Berthet 2009) |
| 23 | Vavalle et al.(Vavalle et al., 2015) | M50-O | - | - | - | x | x | x | Thorax frontal impact tests (Lebarbe et al. 2012) blunt lateral impact to upper extremity (Koh et al. 2005) Abdominal impact tests (Hardy et al. 2001) Pelvis lateral impact tests (Bouquet et al. 1998) |
| 24 | Untaroiu et al.(Untaroiu et al., 2015) | M50-P | x | x | x | x | x | x | Knee joint under valgus bending (Bose et al. 2008) blunt lateral impact to lower extremity (Viano 1989) Lumbar spine test (Rohlmann et al. 2001) Car to Pedestrian Collusion PMHS test (Kerrigan et al. 2007). |
| 25 | Vavalle et al.(Vavalle et al., 2013) | M50-O | - | - | - | x | x | x | Lateral impact tests (Pintar et al. 1997) Lateral impact tests (Cavanaugh et al. 1990) Lateral drop test (Stalnaker et al. 1979) |

* FBM – full body model;* RBM – regional body model;* PLX– pelvic-lower extremity

Note: The head, neck, thorax, abdomen and pelvic-lower-extremities models are validated for M50-O model and no changes were made to these models for pedestrian (M50-P) model except change of posture from seated to standing.

**Supplementary Table S2.** Material properties of the brain used in baseline simulations (Mao et al.(Mao et al., 2013)).

| Component | Density ($kg/m^3$) | Bulk modulus (GPa) | Short-time shear modulus (kPa) | Long-time shear modulus (kPa) | Decay constant $sec^{-1}$ |
|---|---|---|---|---|---|
| Cerebrum gray matter, Cerebellum, Thalamus, Brainstem, Basal ganglia | 1060 | 2.19 | 6.0 | 1.2 | 80 |
| CSF, 3rd Ventricle, Later ventricle | 1040 | 2.19 | 0.5 | 0.1 | 80 |
| Corpus callosum, Cerebrum white matter | 1060 | 2.19 | 7.5 | 1.5 | 80 |

**Supplementary Table S3.** Material properties of the brain used in the parametric study (Chafi et al.(Chafi et al., 2011)). Material properties are based on high strain rate response.

| |
|---|
| Hyperelastic terms |
| Mooney–Rivlin model<br>$K = 2.19\ GPa$ ; $C_{10} = 514.62$ Pa ; $C_{01} = 566.08$ Pa |
| Viscoelastic terms for frequency range of 0.02 Hz–10 MHz |
| Shear modulus:<br>$G_{\infty} = 2160$ Pa ; $G_1 = 156{,}488.3$ kPa ; $G_2 = 326{,}025.8$ kPa; $G_3 = 0.0016$ kPa;<br>$G_4 = 1.2313$ kPa ; $G_5 = 17.583$ kPa; $G_6 = 0.0254$ kPa<br>Decay constant:<br>$\beta_1 = 1.0763 \times 10^9$ sec$^{-1}$; $\beta_2 = 35.8 \times 10^6$ sec$^{-1}$; $\beta_3 = 383.5146 \times 10^3$ sec$^{-1}$; $\beta_4 = 1.0 \times 10^3$ sec$^{-1}$; $\beta_5 = 10$ sec$^{-1}$; $\beta_6 = 3.6533$ sec$^{-1}$ |

**Supplementary Table S4.** Quantitative comparison of reflected pressures from the simulations against the experiments of Skotak et al.(Skotak et al., 2018)

| Comparison case | CORA score<br>experiment (Skotak et al., 2018) vs. simulation |
| --- | --- |
| Reflected overpressure, H1 | 0.754 |
| Reflected overpressure, H2 | 0.523 |
| Reflected overpressure, H3 | 0.815 |
| Reflected overpressure, H4 | 0.755 |
| Reflected overpressure, H5 | 0.724 |

**Supplementary Table S5.** Quantitative comparison of intracranial pressures from the simulations against the experiments of Bir et al.(Bir, 2011)

| Brain pressure | 70 kPa blast intensity | 100 kPa blast intensity |
|---|---|---|
| | CORA score experiment(Bir, 2011) vs. simulation | CORA score experiment(Bir, 2011) vs. simulation |
| Front | 0.228 | 0.584 |
| Occipital | 0.240 | 0.258 |
| Parietal | 0.135 | 0.243 |

**Supplementary Information References:**

Aira, J., Guleyupoglu, B., Jones, D., Koya, B., Davis, M., Gayzik, F.S., 2019. Validated thoracic vertebrae and costovertebral joints increase biofidelity of a human body model in hub impacts. Traffic Inj. Prev. 20, S1–S6. https://doi.org/10.1080/15389588.2019.1638511

Arun, M.W.J.J., Umale, S., Humm, J.R., Yoganandan, N., Hadagali, P., Pintar, F.A., 2016. Evaluation of kinematics and injuries to restrained occupants in far-side crashes using full-scale vehicle and human body models. Traffic Inj. Prev. 17, 116–123. https://doi.org/10.1080/15389588.2016.1197394

Beillas, P., Berthet, F., 2012. Performance Of A 50th Percentile Abdominal Model For Impact: Effects Of Size And Mass. J. Biomech. 45, S83. https://doi.org/10.1016/s0021-9290(12)70084-2

Bir, C., 2011. Measuring Blast-Related Intracranial Pressure Within the Human Head, U.S. Army Medical Research and Materiel Comma, Award number W81XWH-09-1-0498.

Chafi, M.S., Ganpule, S., Gu, L., Chandra, N., 2011. Dynamic response of brain subjected to blast loadings: Influence of frequency ranges. Int. J. Appl. Mech. 3, 803–823. https://doi.org/10.1142/S175882511100124X

Chandra, N., Sundaramurthy, A., Gupta, R.K., 2017. Validation of Laboratory Animal and Surrogate Human Models in Primary Blast Injury Studies. Mil. Med. 182, 105–113. https://doi.org/10.7205/milmed-d-16-00144

Decker, W., Koya, B., Davis, M.L., Gayzik, F.S., 2017. Modular use of human body models of varying levels of complexity: Validation of head kinematics. Traffic Inj. Prev. 18, S155–S160. https://doi.org/10.1080/15389588.2017.1315637

Devane, K., Johnson, D., Gayzik, F.S., 2019. Validation of a simplified human body model in relaxed and braced conditions in low-speed frontal sled tests. Traffic Inj. Prev. https://doi.org/10.1080/15389588.2019.1655733

Fice, J.B., Cronin, D.S., Panzer, M.B., 2011. Cervical spine model to predict capsular ligament response in rear impact. Ann. Biomed. Eng. 39, 2152–2162. https://doi.org/10.1007/s10439-011-0315-4

Gaewsky, J.P., Jones, D.A., Ye, X., Koya, B., McNamara, K.P., Gayzik, F.S., Weaver, A.A., Putnam, J.B., Somers, J.T., Stitzel, J.D., 2019. Modeling Human Volunteers in Multidirectional, Uni-axial Sled Tests Using a Finite Element Human Body Model. Ann. Biomed. Eng. 47, 487–511. https://doi.org/10.1007/s10439-018-02147-3

Ganpule, Daphalapurkar, N.P., Ramesh, K.T., Knutsen, A.K., Pham, D.L., Bayly, P. V., Prince, J.L., 2017. A Three-Dimensional Computational Human Head Model That Captures Live Human Brain Dynamics. J. Neurotrauma 34, 2154–2166. https://doi.org/10.1089/neu.2016.4744

Garimella, H.T., Kraft, R.H., Przekwas, A.J., 2018. Do blast induced skull flexures result in axonal deformation? PLoS One 13, 1–25. https://doi.org/10.1371/journal.pone.0190881

Gayzik, F., Hamilton, C., Tan, J., McNally, C., Duma, S., Klinich, K., Stitzel, J., 2009. A multi-modality image data collection protocol for full body finite element model development. SAE Tech. Pap. 4970. https://doi.org/10.4271/2009-01-2261

Gayzik, F., Moreno, D., Geer, C., Wuertzer, S., Martin, R., Stitzel, J., 2011a. Development of a full body CAD dataset for computational modeling: A multi-modality approach. Ann. Biomed. Eng. 39, 2568–2583.

https://doi.org/10.1007/s10439-011-0359-5

Gayzik, F., Moreno, D., Vavalle, N., Rhyne, A., Stitzel, J., 2011b. Development of the Global Human Body Models Consortium mid-sized male full body model. Inj. Biomech. Res. 39–12.

Giordano, C., Kleiven, S., 2014. Evaluation of Axonal Strain as a Predictor for Mild Traumatic Brain Injuries Using Finite Element Modeling. SAE Tech. Pap. 2014-Novem. https://doi.org/10.4271/2014-22-0002

Hayes, A.R., Vavalle, N.A., Moreno, D.P., Stitzel, J.D., Gayzik, F.S., 2014. Validation of Simulated Chestband Data in Frontal and Lateral Loading Using a Human Body Finite Element Model. Traffic Inj. Prev. 15, 181–186. https://doi.org/10.1080/15389588.2013.799278

Ji, S., Zhao, W., Ford, J.C., Beckwith, J.G., Bolander, R.P., Greenwald, R.M., Flashman, L.A., Paulsen, K.D., McAllister, T.W., 2015. Group-wise evaluation and comparison of white matter fiber strain and maximum principal strain in sports-related concussion. J. Neurotrauma 32, 441–454. https://doi.org/10.1089/neu.2013.3268

Kindig, M., Li, Z., Kent, R., Subit, D., 2015. Effect of intercostal muscle and costovertebral joint material properties on human ribcage stiffness and kinematics. Comput. Methods Biomech. Biomed. Engin. 18, 556–570. https://doi.org/10.1080/10255842.2013.820718

Knutsen, A.K., Gomez, A.D., Gangolli, M., Wang, W.-T., Chan, D., Lu, Y.-C., Christoforou, E., Prince, J.L., Bayly, P. V, Butman, J.A., Pham, D.L., 2020. In vivo estimates of axonal stretch and 3D brain deformation during mild head impact.

Brain Multiphysics 1, 100015. https://doi.org/https://doi.org/10.1016/j.brain.2020.100015

Levine, J., Dionne, J.P., Bueley, D., Makris, A., 2014. Blast and Impact Induced Linear and Rotational Head Acceleration. Pass 2014.

Li, Z., Kindig, M.W., Subit, D., Kent, R.W., 2010. Influence of mesh density, cortical thickness and material properties on human rib fracture prediction. Med. Eng. Phys. 32, 998–1008. https://doi.org/10.1016/j.medengphy.2010.06.015

Mao, H., Zhang, L., Jiang, B., Genthikatti, V. V., Jin, X., Zhu, F., Makwana, R., Gill, A., Jandir, G., Singh, A., Yang, K.H., 2013. Development of a finite element human head model partially validated with thirty five experimental cases. J. Biomech. Eng. 135. https://doi.org/10.1115/1.4025101

Miller, L.E., Urban, J.E., Stitzel, J.D., 2017. Validation performance comparison for finite element models of the human brain. Comput. Methods Biomech. Biomed. Engin. 20, 1273–1288. https://doi.org/10.1080/10255842.2017.1340462

Panzer, M.B., Fice, J.B., Cronin, D.S., 2011. Cervical spine response in frontal crash. Med. Eng. Phys. 33, 1147–1159. https://doi.org/10.1016/j.medengphy.2011.05.004

Park, G., Kim, T., Panzer, M.B., Crandall, J.R., 2016. Validation of Shoulder Response of Human Body Finite-Element Model (GHBMC) Under Whole Body Lateral Impact Condition. Ann. Biomed. Eng. 44, 2558–2576. https://doi.org/10.1007/s10439-015-1546-6

Poulard, D., Kent, R.W., Kindig, M., Li, Z.P., Subit, D., 2015. Thoracic response targets for a computational model: A hierarchical approach to assess the biofidelity of a 50th-percentile occupant male finite element model. J. Mech. Behav. Biomed.

Mater. 45, 45–64. https://doi.org/10.1016/j.jmbbm.2015.01.017

Schap, J.M., Koya, B., Gayzik, F.S., 2019. Objective Evaluation of Whole Body Kinematics in a Simulated, Restrained Frontal Impact. Ann. Biomed. Eng. 47, 512–523. https://doi.org/10.1007/s10439-018-02180-2

Shin, J., Untaroiu, C.D., 2013. Biomechanical and injury response of human foot and ankle under complex loading. J. Biomech. Eng. 135. https://doi.org/10.1115/1.4025108

Skotak, M., Alay, E., Zheng, J.Q., Halls, V., Chandra, N., 2018. Effective testing of personal protective equipment in blast loading conditions in shock tube: Comparison of three different testing locations. PLoS One 13, 1–19. https://doi.org/10.1371/journal.pone.0198968

Somasundaram, K., Zhang, L., Sherman, D., Begeman, P., Lyu, D., Cavanaugh, J.M., 2019. Evaluating thoracolumbar spine response during simulated underbody blast impact using a total human body finite element model. J. Mech. Behav. Biomed. Mater. 100, 103398. https://doi.org/10.1016/j.jmbbm.2019.103398

Untaroiu, C.D., Tech, V., Schap, J., Putnam, J.B., Tech, V., Gayzik, F.S., Davis, M.L., 2015. Detc2015-47781 Pedestrian Finite Element Model. Proc. ASME 2015 Int. Des. Eng. Tech. Conf. Comput. Inf. Eng. Conf. 1–7.

Untaroiu, C.D., Yue, N., Shin, J., 2013. A finite element model of the lower limb for simulating automotive impacts. Ann. Biomed. Eng. 41, 513–526. https://doi.org/10.1007/s10439-012-0687-0

Vavalle, N.A., Davis, M.L., Stitzel, J.D., Gayzik, F.S., 2015. Quantitative Validation of a Human Body Finite Element Model Using Rigid Body Impacts. Ann. Biomed. Eng.

43, 2163–2174. https://doi.org/10.1007/s10439-015-1286-7

Vavalle, N.A., Moreno, D.P., Rhyne, A.C., Stitzel, J.D., Gayzik, F.S., 2013. Lateral impact validation of a geometrically accurate full body finite element model for blunt injury prediction. Ann. Biomed. Eng. 41, 497–512. https://doi.org/10.1007/s10439-012-0684-3

Weaver, C.M., Baker, A.M., Davis, M.L., Miller, A.N., Stitzel, J.D., 2018. Finite Element-Based Pelvic Injury Metric Creation and Validation in Lateral Impact for a Human Body Model. J. Biomech. Eng. Asme 140. https://doi.org/10.1115/1.4039393

Xiao, S., Yang, J., Huang, J., Crandall, J.R., 2017. Development Of A Belted Occupant Fe Model For Prediction Of Chest Injury Risk Based On Stress And Strain Analysis. J. Mech. Med. Biol. 17. https://doi.org/10.1142/s0219519417500609

Yue, N., Untaroiu, C.D., 2014. A Numerical Investigation on the Variation in Hip Injury Tolerance With Occupant Posture During Frontal Collisions. Traffic Inj. Prev. 15, 513–522. https://doi.org/10.1080/15389588.2013.840884

Zhao, W., Ji, S., 2018. White Matter Anisotropy for Impact Simulation and Response Sampling in Traumatic Brain Injury. J. Neurotrauma 36, 250–263. https://doi.org/10.1089/neu.2018.5634

Zheng, J.J., Tang, L., Hu, J.W., 2018. A Numerical Investigation of Risk Factors Affecting Lumbar Spine Injuries Using a Detailed Lumbar Model. Appl. Bionics Biomech. 2018. https://doi.org/10.1155/2018/8626102
